# Repair of Mismatched Templates during Rad51-dependent Break-Induced Replication

**DOI:** 10.1101/2022.01.30.478419

**Authors:** Jihyun Choi, Danielle N. Gallagher, Kevin Li, Gabriel Bronk, James E. Haber

## Abstract

Using budding yeast, we have studied Rad51-dependent break-induced replication (BIR), where the invading 3’ end of a site-specific double-strand break (DSB) and a donor template share 108 bp of homology that can be easily altered. When every 10^th^ base is mismatched between donor and recipient, BIR is 44% efficient compared to fully homologous sequences; but BIR still occurs about 10% when every 6^th^ base is mismatched. Here we explore the tolerance of mismatches in more detail, by examining donor templates that each carry 10 mismatches, but where they are clustered with spacings of every 6^th^ bp. These different arrangements of uneven mismatch distribution were in general less efficient in recombination as templates with evenly distributed mismatches. A donor with all 10 mismatches clustered every 6^th^ base at the 3’ invading end of the DSB was not impaired compared to the case where mismatches were clustered at the 5’ end. These data suggest that the efficiency of strand invasion is principally dictated by thermodynamic considerations, i.e., by the total number of base pairs that can be formed; but sequence-specific factors are also important. Mismatches in the donor template are incorporated into the BIR product in a strongly polar fashion up to ~40 nucleotides from the 3’ end. Mismatch incorporation depends on the 3’ → 5’ proofreading exonuclease activity of DNA polymerase δ, with little contribution from Msh2/Mlh1 mismatch repair proteins. Surprisingly, the probability of a mismatch 27 nt from the 3’ end being replaced by donor sequence was the same whether the preceding 26 nucleotides were mismatched every 6^th^ base or fully homologous. These data suggest that DNA polymerase δ “chews back” the 3’ end of the invading strand without any mismatch-dependent cues from the strand invasion structure.

**Author Summary:** DNA double-strand breaks (DSBs) are the most lethal forms of DNA damage and inaccurate repair of these breaks presents a serious threat to genomic integrity and cell viability. Break-induced replication (BIR) is a homologous recombination pathway that results in a nonreciprocal translocation of chromosome ends. We used budding yeast *Saccharomyces cerevisiae* to investigate Rad51-mediated BIR, where the invading 3’ end of the DSB and a donor template share 108 bp of homology. We examined the tolerance of differently distributed mismatches on a homologous donor template and found that BIR efficiency was the same whether the mismatches were clustered at the 3’ invading end or at the 5’ end. We confirmed that mismatches are incorporated into the BIR product in a strongly polar fashion as far as about 40 nucleotides from the 3’ end. We conclude that the proofreading activity of DNA polymerase δ “chews back” the 3’ end of the invading strand even when the sequences removed have no mismatches for the first 26 nucleotides. These observations enrich our understanding of the details of Rad51-mediated strand invasion and provide insight into the mechanism of the 3’ to 5’ proofreading activity of DNA polymerase during homologous recombination.

## Introduction

DNA double-strand breaks (DSBs) are the most toxic lesions that can occur in DNA, and failure to repair these breaks can result in genome instability. Eukaryotes have evolved two major types of DNA repair mechanisms to deal with DSBs: non-homologous end joining (NHEJ) and homologous recombination (HR). In both “classic” NHEJ and in microhomology-mediated end-joining, broken ends are ligated back together, often using a small amount of microhomology at the junction, frequently resulting in small insertions and deletions [1,2,3,4]. HR by gene conversion relies on the resected ends of a DSB searching for and copying an intact homologous sequence that serves as a template to accurately repair the lesion [5]. Two other HR repair pathways - single-strand annealing (SSA) and break-induced replication (BIR) – also rely on pairing with homologous sequences but often result in chromosome alterations [5].

BIR repairs DSBs that share only one end of homology with a donor sequence; such events allow the extension of eroded telomeres and the re-initiation of DNA replication at stalled and broken replication forks [6,7]. BIR is initiated by 5’ → 3’ resection of a broken DNA end to generate a 3’ single-stranded DNA (ssDNA) tail (Fig. S1A). Initially, replication protein complex A (RPA) coats the ssDNA tails but is then displaced by the recombination protein, Rad51 [8,9]. Each monomer of Rad51 and its bacterial homolog, RecA, binds 3 nucleotides of ssDNA to form a nucleoprotein filament that catalyzes base-pairing and strand invasion between the Rad51-coated ssDNA end of DSB and a homologous double-stranded DNA (dsDNA) donor [10,11,12,13]. Strand invasion and the formation of a displacement loop (D-loop) enables DNA polymerase δ to prime DNA synthesis and extend the 3’ end of invading strand to the end of the chromosome [14,15,16]. In many respects, BIR events studied in budding yeast resemble mammalian alternative lengthening of telomeres (ALT) [7,17,18]. BIR also appears to be critical for DNA synthesis that occurs very late in the cell cycle as cells enter mitosis (MiDAS) [7,19,20,21]. Both in yeast and in mammalian cells, the nonessential Pol32 (POLD3) subunit of DNA polymerase δ is required for BIR but not for normal DNA replication [22,23,24].

Precisely how Rad51 performs strand exchange remains a subject of great interest [25,26,27,28]. In budding yeast, efficient recombination by gene conversion (where both ends of the DSB interact with a donor template) requires a homologous region of about 70 bp [27,29]. Rad51-dependent BIR also has similar homology length requirements. For example, in an intrachromosomal BIR assay (Fig. 1) in which a site-specific DSB is induced by the expression of HO endonuclease, approximately 14% of cells successfully repaired a DSB when the homologous region is 108 bp; but repair dropped to 1% with a 54-bp template and to 0.02% when there were only 26 bp [24]. Repair is also reduced by the presence of heterologies (the presence of non-complementary bases between donor and recipient sequences), although even when every 6^th^ base was mismatched, BIR occurred at 10% of the rate when the templates were identical [24]. Repair efficiency was significantly reduced but still significant when every 6^th^ base was mismatched. These *in vivo* results contrast with *in vitro* single-molecule studies using very short substrates that found a minimum of at least 8 consecutive bases must be paired for a stable initial strand exchange mediated by RecA or Rad51 [10]. Another *in vitro* study concluded that 8 contiguous base pairs are required to tolerate up to a single mismatch in RecA-mediated strand exchange [30,31]. The differences among these studies may reflect the longer (108-bp) substrates used in our *in vivo* studies or the agency of other recombination proteins not presented in the *in vitro* studies.

**Figure 1.**
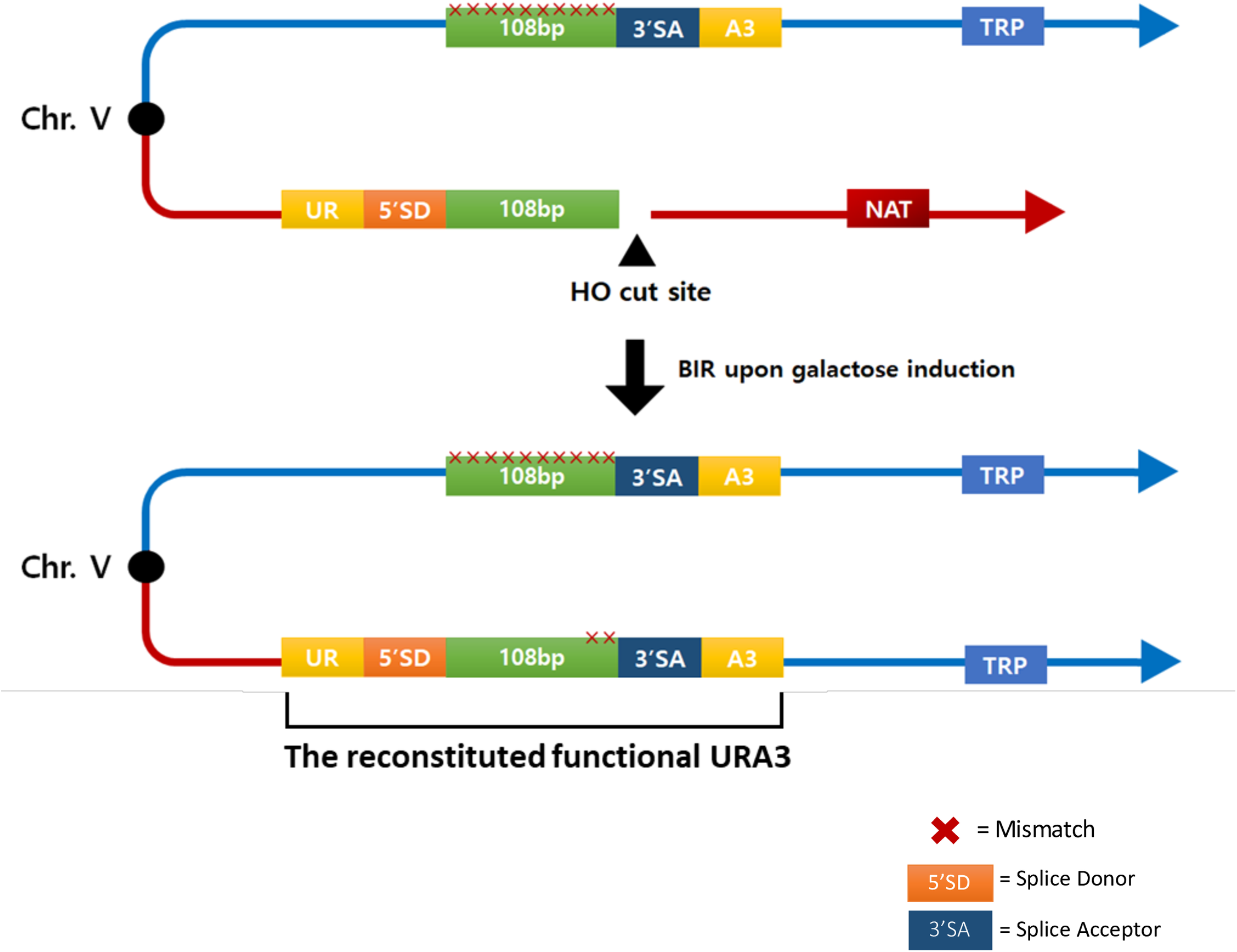
BIR-dependent formation of a functional Ura3^+^ recombinant. The recipient sequence shares the 108-bp region of homology and contain the 5’ sequences from the *URA3* gene (UR), the splice donor site (5’ SD) of an artificial intron and the HO cut site. 108-bp donor sequences containing different mismatch distributions were assembled into a plasmid containing the 3’ sequences from the *URA3* gene (A3), the 3’ splice-acceptor (3’ SA) of the intron and the *TRP1* auxotrophic maker. A DSB was created using the galactose-inducible HO endonuclease. This break is repaired by BIR using the donor sequences that share 108-bp of homology located on the opposite arm of chromosome V. Once BIR is complete, a functional intron is formed, and yeast become Ura3^+^ recombinants.

Here, we extended our studies of mismatch tolerance by examining different templates with the same number of mismatches but distributed unevenly. We wished to determine if these different distributions would be treated equivalently, which would be the case if strand invasion in BIR were principally governed by thermodynamic considerations, in which the total number of base pairs that can be formed dictates repair efficiency. We show that these different arrangements were in general less efficient than the evenly distributed mismatches, but exhibit significant differences that depend on the precise location of the mismatches. We show that this discrimination does not appear to be monitored by the Msh2-dependent mismatch repair system.

A second important aspect of our analysis of repair involving mismatched substrates came from analyzing the assimilation of heterologies into the BIR product. Mismatches close to the 3’ end of the invading strand are almost invariably replaced by the template sequence, but there is a steep decline in their incorporation, so that by about 40 bp from the 3’ end, incorporation of the template sequence is rare [24]. The assimilation of the template sequence is dependent on the 3’ to 5’ proofreading activity of DNA polymerase δ, which presumably chews back the 3’ end of the invading strand before initiating copying of the donor template (Fig. S1B). There was little effect when Msh2/Mlh1-dependent mismatch repair was ablated, although the extent of mismatch assimilation was shortened [24]. Here, we make the surprising discovery that 3’ to 5’ removal of the 3’ end of the strand-invading DNA is evident even when the first 26 nucleotides are completely homologous to the template, suggesting that this resection is not provoked by a nearby mismatch.

## Results

To study the effect of different distributions of mismatches on BIR repair efficiency we used the assay shown in Fig. 1. A galactose-inducible site-specific DSB is created by HO endonuclease at a site just distal to the 5’ end of the *URA3* gene (UR) joined to an mRNA splicing donor sequence (SD) [24]. Only one end of the DSB shares homology with the donor, which in this case is located on the opposite arm of the same chromosome, about 30 kb from the telomere. The donor consists of a 108-bp region of homology such that the DSB end is perfectly matched to the donor (i.e., there are no additional nonhomologous sequences at the 3’ end). The 108-bp homologous segment is adjacent to 3’ splice acceptor (SA) site, followed by the 3’ end of the *URA3* gene (A3). Thus, BIR results in a nonreciprocal translocation producing an intron-containing, intact *URA3* gene so that cells can grow in the absence of uracil.

We confirmed our previous results [24] that strain yRA280 (with every 10^th^ base pair mismatched) and yRA321 (with every 6^th^ base pair mismatched) had reduced, but significant levels of repair (Fig. 2B) compared to yRA253 (a fully homologous donor). BIR was reduced to about 44% in yRA280 compared to yRA253, and still occurred about 9% in yRA321 (Fig. 2B). We then created six new strains, each of which had 10 mismatches, but arranged so that they were clustered every 6 bases apart (Fig.2A). The mismatches included both transversion and transition mismatches (Table 2).

**Figure 2.**
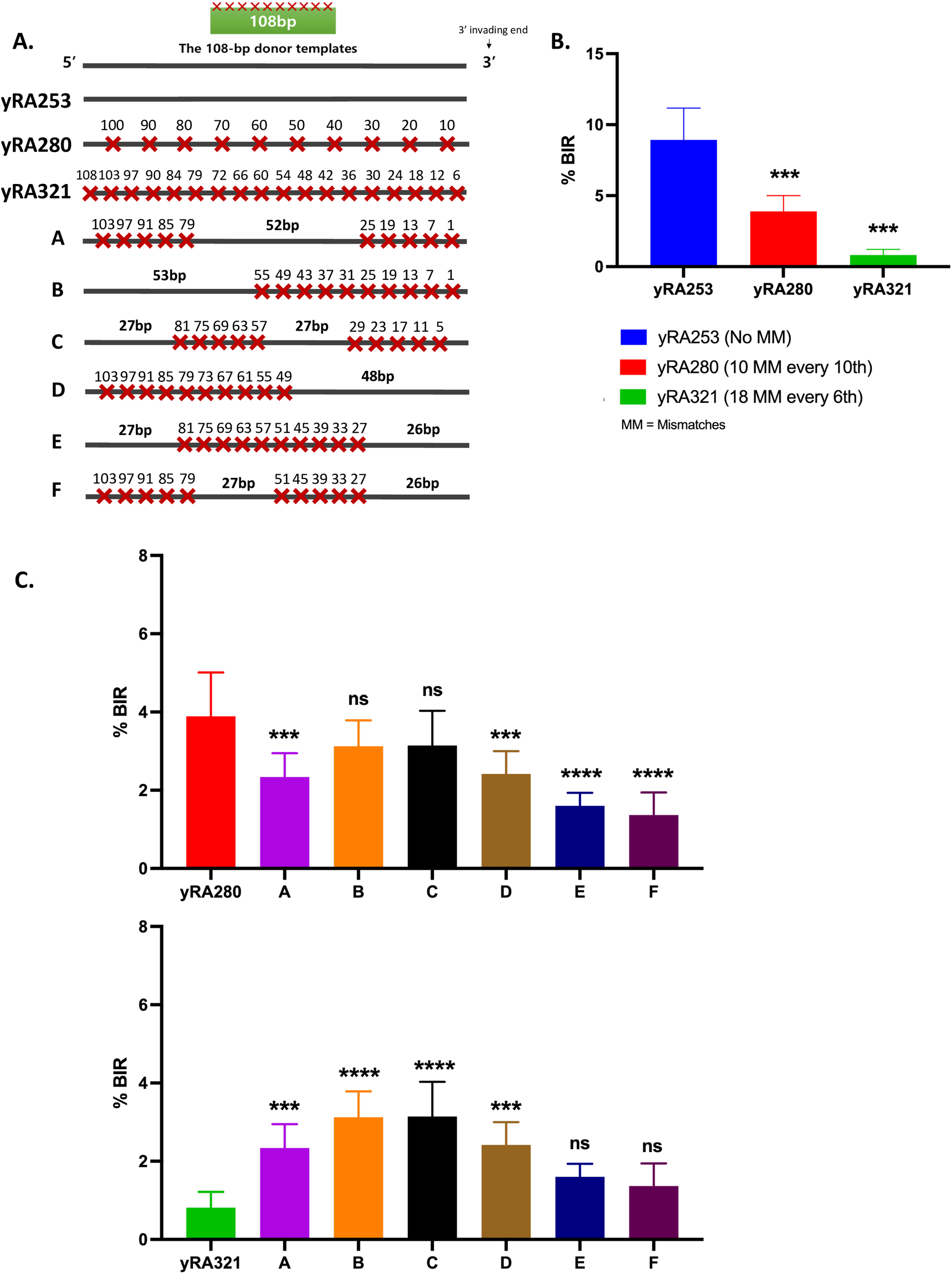
The effect of different distributions of mismatches on a 108-bp donor template on the efficiency of Rad51-dependent BIR. **A.** 108-bp donors with even mismatch distribution: yRA253 (no mismatch), yRA280 (10 mismatches every 10th) and yRA321 (18 mismatches every 6th). The set of donors with even mismatch distribution was compared with a set of divergent 108-bp donor templates containing a total of 10 mismatches that are distributed unevenly throughout the template. The spacing between the clustered mismatches are every 6th bp. Specific sequences are shown in Table 2. **B.** Percent BIR efficiency of 108-bp donors with perfect homology and even mismatch distribution. Welch’s t-test was used to determine the p-value, **p ≤ 0.001, ns = not significant. Error bars indicate standard deviation. A minimum of three measurements were performed. **C.** Percent BIR efficiency of 108-bp donors with even and uneven distribution. The set of donors with uneven mismatch distribution was compared to yRA280 which contains the same mismatch density as all unevenly mismatched donors and to yRA321 which has the same 6th bp spacing between clustered mismatches. Significance determined using a Dunnett’s method (GraphPad Prism 9). Error bars refer to standard deviation. ** p<0.001, ns = not significant.

The efficiency of BIR for the substrate with all mismatches clustered at the 3’ invading end of the DSB (Fig. 2C, donor template B) was similar to the reference yRA280, and was surprisingly more efficient than with a substrate with a 48 bp of perfect homology at the 3’ end (Fig. 2C, donor template D). Both template B and D were significantly different from reference strain yRA321, with every 6^th^ bp mismatched and 18 total mismatches (Fig. 2C). Thus, although the 3’ end must be synapsed with the donor to allow DNA polymerase to initiate new DNA synthesis at the 3’ end, the efficiency of repair was not more impaired than with the mismatches all at the opposite end. These conclusions are generally supported by the results using other mismatch spacings (Fig. 2A); but other substrates with 26 bp of perfect homology at the 3’ end (Fig. 2C, donor templates E and F) yielded a significantly lower rate of successful recombination when compared to yRA280 (Fig. 2C). The differences among these templates cannot be attributed to a difference in thermal stability of base-pairing in the 108-bp region as measured by the calculated melting temperature (Tm) between complementary 108-nt DNA strands (Table 2 and Fig. 3A). Interestingly, the correlation between BIR efficiency and Tm, seen as the increasing frequency as a function of Tm, was nearly the same among the 6 unevenly-spaced templates (dotted line, Fig. 3A) as for the evenly-spaced cases, but the correlation was shifted to a lower average value (see Discussion). We note also that we did not find any difference in the thermal stability of possible secondary structures that could be formed by the different 108-nt regions, viewed as single-strand sequences (see Methods).

**Figure 3.**
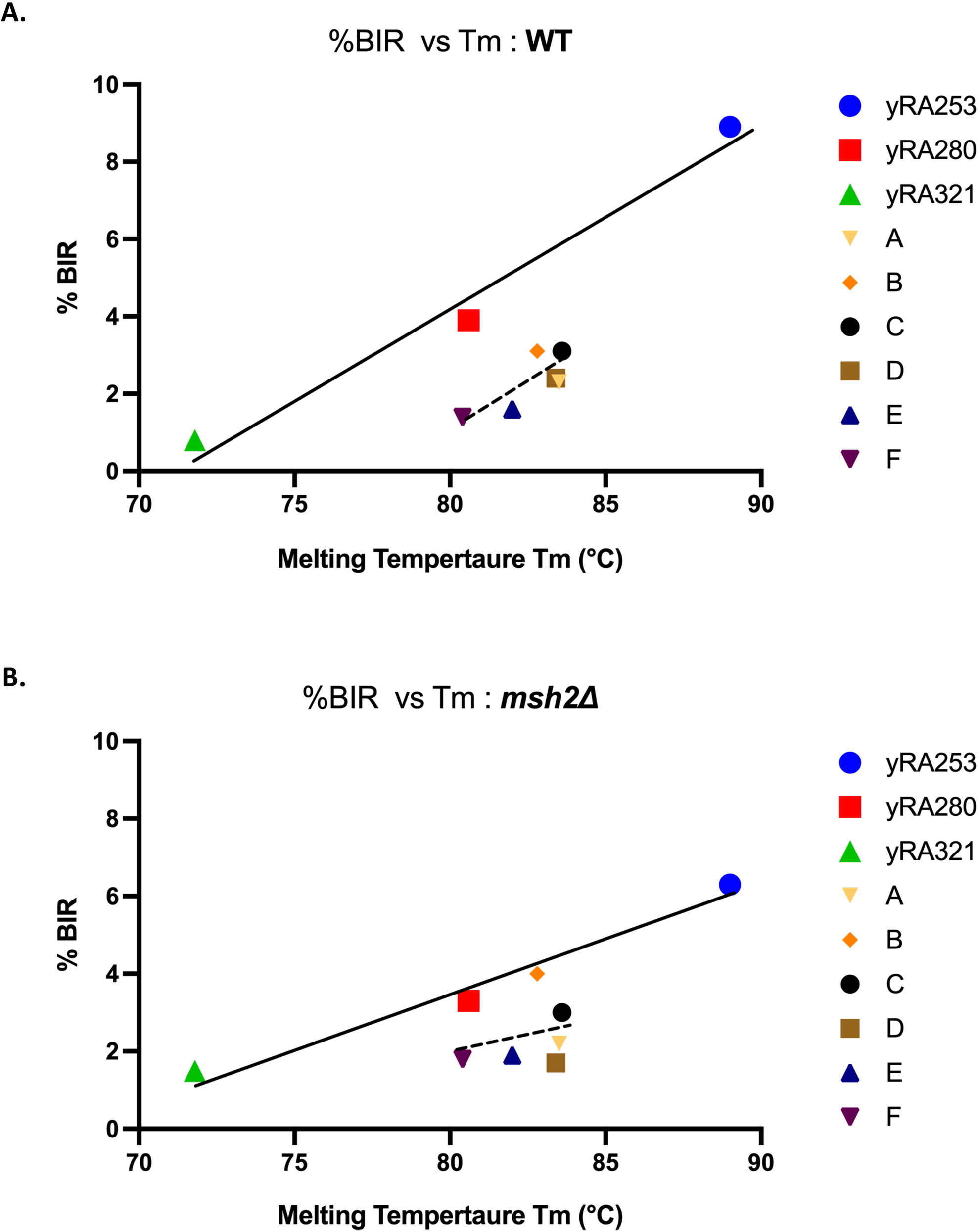
Rad51-mediated strand annealing is less efficient when the mismatches are spaced every 6bp apart. A. Percent BIR of both evenly and unevenly mismatched donor templates are plotted against their melting temperatures Tm (°C). The melting temperature of single-stranded DNA of each donor template, synapsed with the complementary single-strand DNA of recipient sequence (Table 2), was calculated by the method of Markham and Zucker [37] http://www.unafold.org/Dinamelt/applications/two-state-melting-hybridization.php. A least-squares line (solid black line) was determined for the three samples with 0, evenly-spaced every 10^th^ and evenly-spaced every 6^th^ bp mismatches (yRA253, yRA280, and yRA321). A least-squares line for the six donor templates with 10 uneven mismatch distributions was plotted separately (black dotted line). B. Similar comparisons were made for strains deleted for *MSH2*.

As we noted above, the 3’ end of the DSB is matched to the 108-bp template, with no nonhomologous 3’ extension. Our previous studies have shown that even a short 3’ extension significantly reduces BIR repair, attributable to the role of the Msh2 mismatch repair protein in triggering heteroduplex rejection [24]. We created a set of *msh2*Δ derivatives of the set of templates and found that the differences between the evenly-spaced mismatches and the uneven cases were not suppressed (Fig. 3 and Fig. S2). However, the efficiency of BIR was generally reduced relative to wild type, even for the case of no mismatches and the differences between yRA280 and yRA321 was no longer statistically significant (Fig. S2).

### Assimilation of mismatches into BIR products reveals the role of DNA polymerase δ

Once heteroduplex DNA forms by strand invasion, mismatch correction may lead to the incorporation of donor sequences into the BIR product. However, the assimilation of heterologies in BIR does not proceed through the general Msh2/Mlh1-dependent mismatch repair system [24], but instead is dependent on the 3’ to 5’ exonuclease activity of DNA polymerase δ, which chews away 3’ end of the invading strand and then copies the donor sequence [24]. Thus, mismatches very close to the 3’ end of the invading DNA are very frequently replaced by the donor allele. There is a steep drop in assimilation, extending 40-50 nt from the DSB end, after which there was little or no incorporation of the mismatches (Fig. 4E) [24].

**Figure 4.**
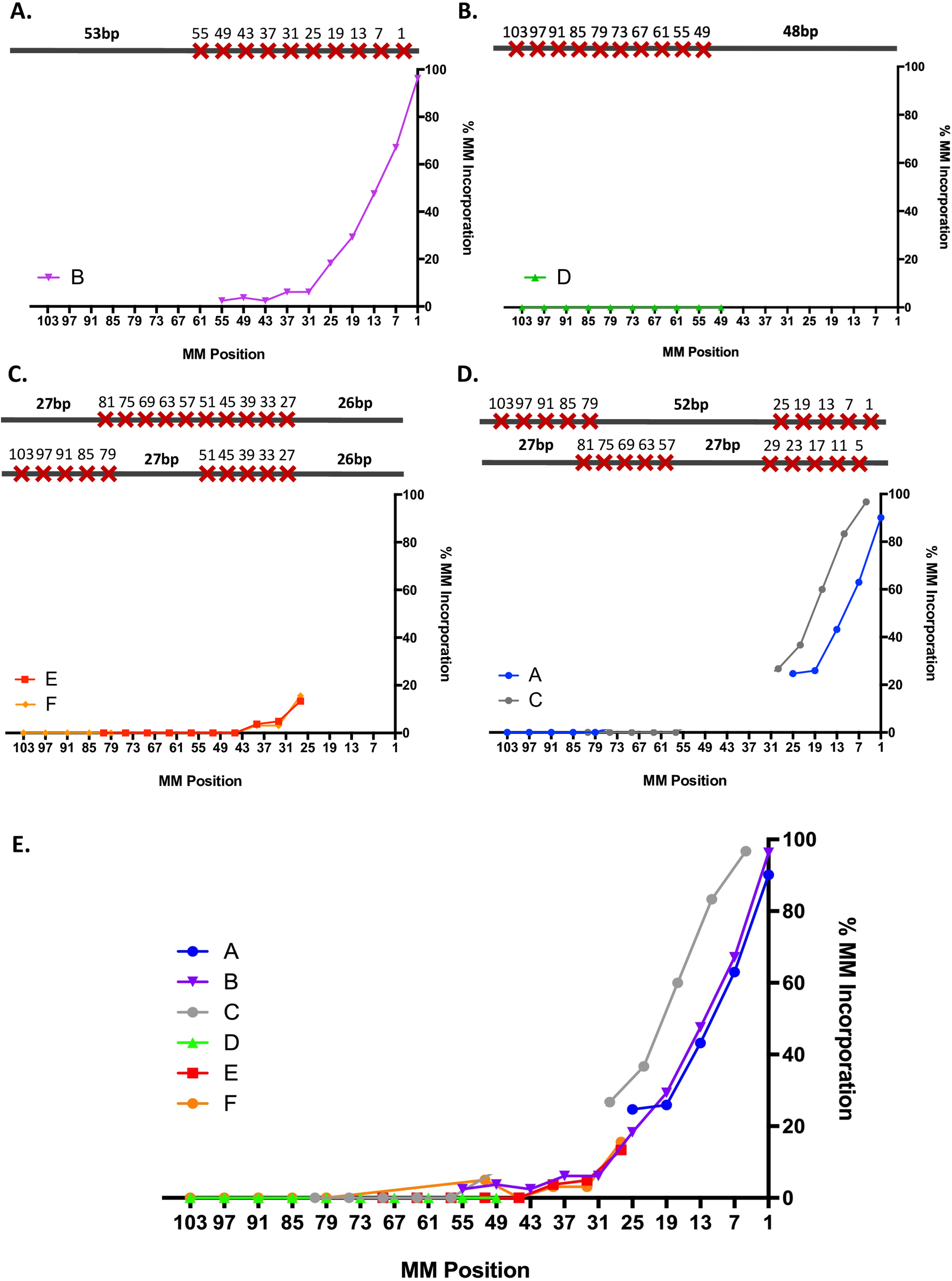
DNA Pol δ still “backs up” to perform 3’ → 5’ proofreading activity to incorporate mismatches even if there are no mismatches closer to the 3’ end of DSB. **A-D.** Percent mismatch incorporation of individual donor templates containing 10 mismatches distributed unevenly throughout the 108-bp donor template. **E.** Composite of % mismatch corrections in 6 different templates with different distributions of 10 mismatches.

To determine the extent of incorporating mismatches into the final product, we sequenced approximately 50 BIR products from each of the templates shown in Fig. 2A. In each case, mismatches were incorporated into the BIR product with the same strong polarity seen with the evenly spaced mismatches [24]. Mismatch assimilation was seen as far as 55 nucleotides from the 3’ invading end (Fig. 4A, donor template B). For template D, containing 10 mismatches clustered near the 5’ end and with 48 bp perfect homology near the 3’ end of the break, none of the 10 mismatches were incorporated (Fig. 4B, template D). However, for all the constructs in which there were mismatches in the first 48bp, the pattern of incorporation was the same (Fig. 4E). Surprisingly, in donor templates E and F (Fig. 4C), the degree of incorporation of mismatches at positions 27, 33 and 39 bp from the invading end was indistinguishable from the correction of these same sites when all the mismatches are present at the 3’ end (Fig. 4A, donor template B). These data suggest that 3’ to 5’ exonuclease activity of DNA polymerase δ removes the 3’ end of the invading strand to incorporate mismatches beyond 26 bp even when this region lacks any mismatches.

To confirm that assimilation of mismatches was independent of the Msh2/Mlh1-dependent mismatch repair system, we examined repair in the two reference strains, yRA280 and yRA321, as well as in donor template A (Fig. 5A). In each case, cells lacking *MLH1* or *MSH2* still can extend and correct mismatches >30 bp from the 3’ end of the DSB (Fig. 5A). There are no consistent differences between wild type, *msh2*Δ and *mlh1*Δ strains; the differences in assimilation most likely reflect the limited number of sequences used to measure mismatch assimilation.

**Figure 5.**
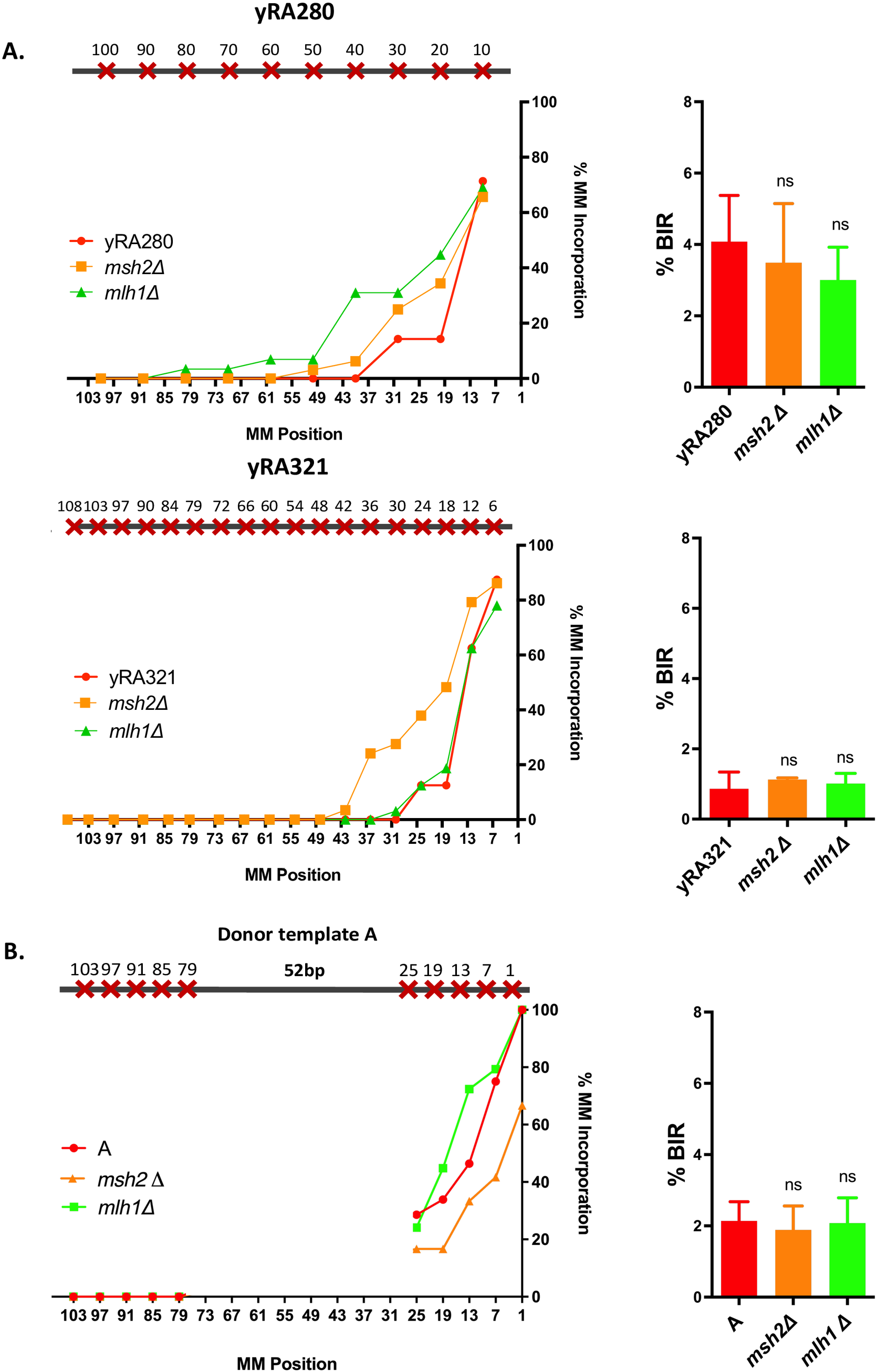
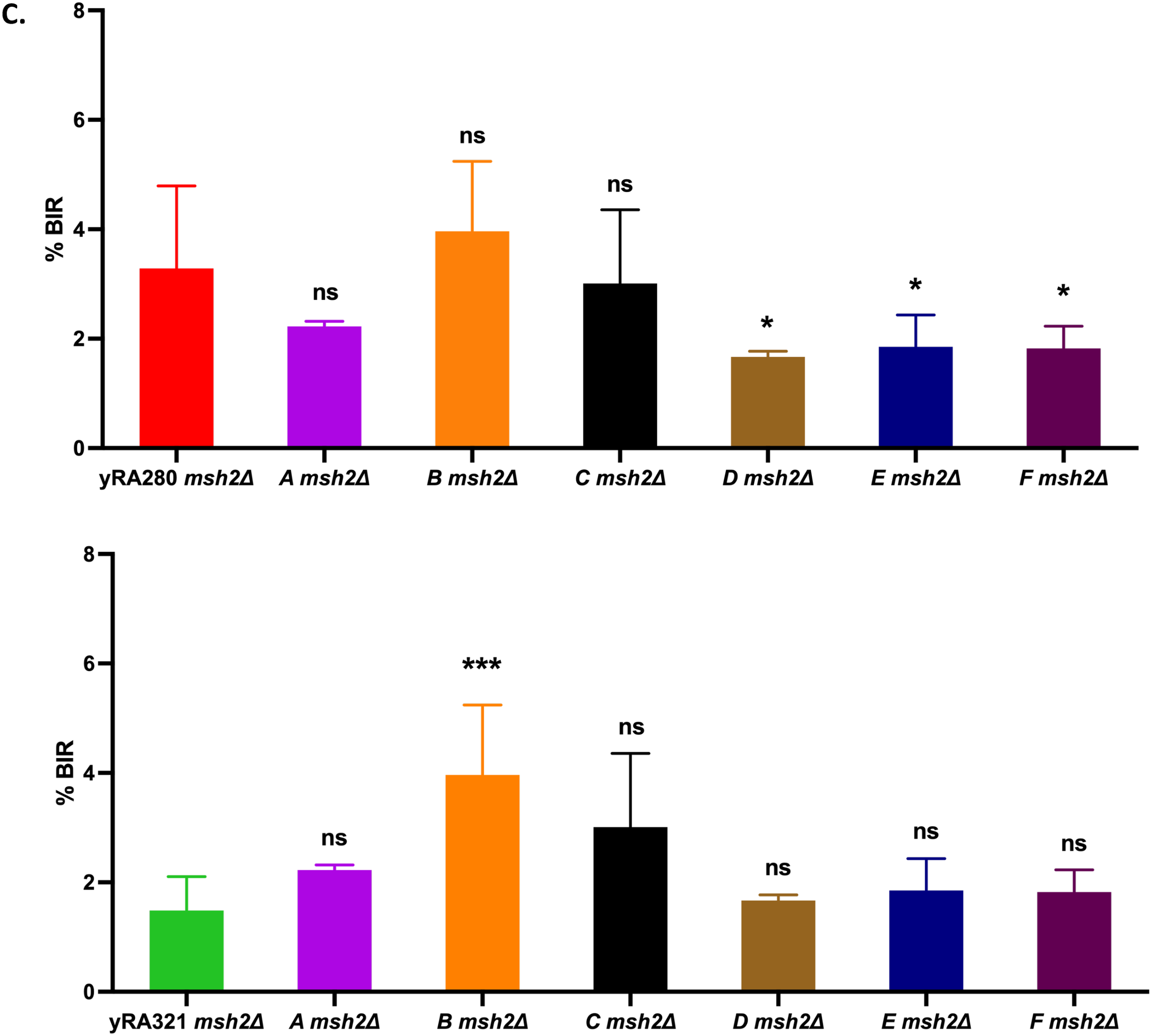
The mismatch assimilation of the template sequence does not proceed through Msh2/Mlh1-dependent mismatch repair. **A.** The effect of deleting mismatch repair gene *MSH2* or *MLH1* on mismatch incorporation pattern and BIR efficiency for strains yRA280 and yRA321. **B.** The effect of deleting mismatch repair gene *MSH2* or *MLH1* on mismatch incorporation pattern and BIR efficiency for donor template A. Welch’s t-test was used to determine the p-value. Error bars refer to standard error of the mean. A minimum of at least three measurements were performed **C.** The effect of deleting mismatch repair gene *MSH2* on BIR efficiency for a set of divergent 108-bp donor templates with uneven distribution. Significance determined using a Dunnett’s method for multiple comparisons. Error bars refer to standard error of the mean.

## Discussion

Although properties of budding yeast Rad51 have been well-studied *in vitro* [11,16,24,33], the ability of Rad51 to search and strand-invade mismatched sequences *in vivo* is still not fully understood. Here we have examined Rad51-dependent BIR where the 3’ invading end of DSB and its donor templates share 108-bp homology, each carrying 10 mismatches, but arranged in several different ways with a spacing of every 6^th^ bp. Donor templates that clustered their mismatches near the 3’ end, where strand invasion may begin (or at least must be annealed before repair DNA synthesis can be initiated) were not impaired in their repair compared to those with mismatches clustered at the 5’ end or in other arrangements. Overall, these results demonstrate that the efficiency of repair is not strongly influenced by perfect homology at the 3’ end and further suggest that the success of repair is primarily a function of the total number of base pairs that can be formed over the entire region. However, none of the arrangements of 10 mismatches with every 6^th^ base spacing was as efficient as a template with 10 mismatches, spaced every 10 base pairs, although in two cases the difference was not statistically significant. If the regions with many mismatches were not base-paired at all, we would expect to see BIR efficiencies comparable to the length of the remaining ~ 54 bases in perfectly-matched sub-regions; however, we have previously shown that a template that had 54 perfectly matched base pairs (i.e. equivalent to the segments of the templates with no mismatches) yielded a BIR rate less than 7% of a 108-bp region [24], whereas the efficiencies of the 6 templates we used were about 26% as successful (and where every 10^th^ was 44%). So the regions containing mismatches every 6^th^ nucleotide must play a role in base-pairing, but there is a significant reduction in efficiency. This penalty is less than the very strong barrier seen from *in vitro* strand exchange studies [10,30,31]. The ability of Rad51 *in vivo* to deal with these highly mismatched regions could be explained by the presence of recombination factors *in vivo* that are not present *in vitro*. As shown in Fig. 3, the 6 templates with uneven mismatch spacing obey a similar temperature-dependence as the evenly-spaced controls; but the success of BIR is reduced (e.g. compare yRA280 and template F, which have almost the same Tm). We have not identified any sequence-specific or other properties of these sequences that can account for the reduced success of BIR, either in initiating recombination or in a later step. Further study will be needed to assess the relation of substrates with different Tm but the same number of base-pairs.

In these assays in which the 3’ end does not have a nonhomologous extension, Msh2 does not provoke a reduction in repair through heteroduplex rejection [24]. Indeed, deleting Msh2 led to an overall reduction in repair efficiency for both the evenly- and unevenly-spaced templates so that any differences became statistically insignificant, but a similar correlation with Tm persists.

The second finding we draw from these experiments concerns the action of DNA polymerase and its 3’ to 5’ proofreading activity. The pattern of mismatch incorporation into the BIR product was the same at interior positions whether or not they were preceded by mismatches nearer the 3’ end; that is, DNA Pol δ still backs up with 3’ → 5’ exonuclease activity to incorporate mismatches even when the first 26 positions are completely homologous. Thus, proofreading at these interior positions is not dependent on any prior mismatch cues and suggests that Pol δ intrinsically backs up on DNA templates even when they are fully homologous. This activity is far more extensive than its apparent action in removing a single mismatched base during DNA replication and is more comparable to the ability of Pol δ to excise a 3’-ended nonhomologous segment at the end of a DSB being repaired by gene conversion [32]; however, the pattern of mismatch assimilation is not compatible with an intermediate in which an unpaired mismatched end would be excised as a 3’ flap, and the incorporation of mismatches would be the same at each position. Deletion of *MSH2* or *MLH1* had no significant effect on the repair outcome.

We note that in meiosis, some gene conversion events appear to have involved a gap-repair near the DSB [33,34,35]. However, one would find a similar pattern if Pol δ were to resect the invading strand in a similar fashion during meiotic DNA repair.

## Methods

### Yeast strains

yRA strains used in these experiments have the haploid S288c background (*ho mat*Δ::hisG *hml*Δ::hisG *hmr*Δ::*ADE3 ura3*Δ-*851 trp1*Δ-63 *leu2*Δ::KAN *ade3*::*GAL10::HO*), in which the site-specific HO endonuclease is under control of a galactose-inducible promoter [36]. Strain yRA111 contains the recipient sequences composed of the 5’ sequences of *URA3* gene (UR), an artificially inserted split-intron with the splice donor site (5’ SD) and the HO recognition site (HOcs), located at the *CAN1* locus in a non-essential terminal region of chromosome V [24]. Divergent donor sequences containing uneven distribution of mismatches were designed and ordered as synthetic gBlocks gene fragments from IDT and assembled into pRS314-based (*CEN4*, *TRP1*) plasmid containing the 3’ splice-acceptor (3’ SA) of the intron, the 3’ sequence from the *URA3* gene (A3) and the *TRP1* marker (called bRA29 plasmid in this study) using *in vivo* recombination and plasmid rescue from a yeast host strain as described in Anand et al. [24]. The donor cassettes were PCR-amplified from plasmid bRA29 and integrated at the *FAU1* locus, about 30 kb from the right end of on chromosome V. Further information for strains, recipients (donors) and plasmids are found in Tables 1, 2, 3 and 4.

**Table 1.**
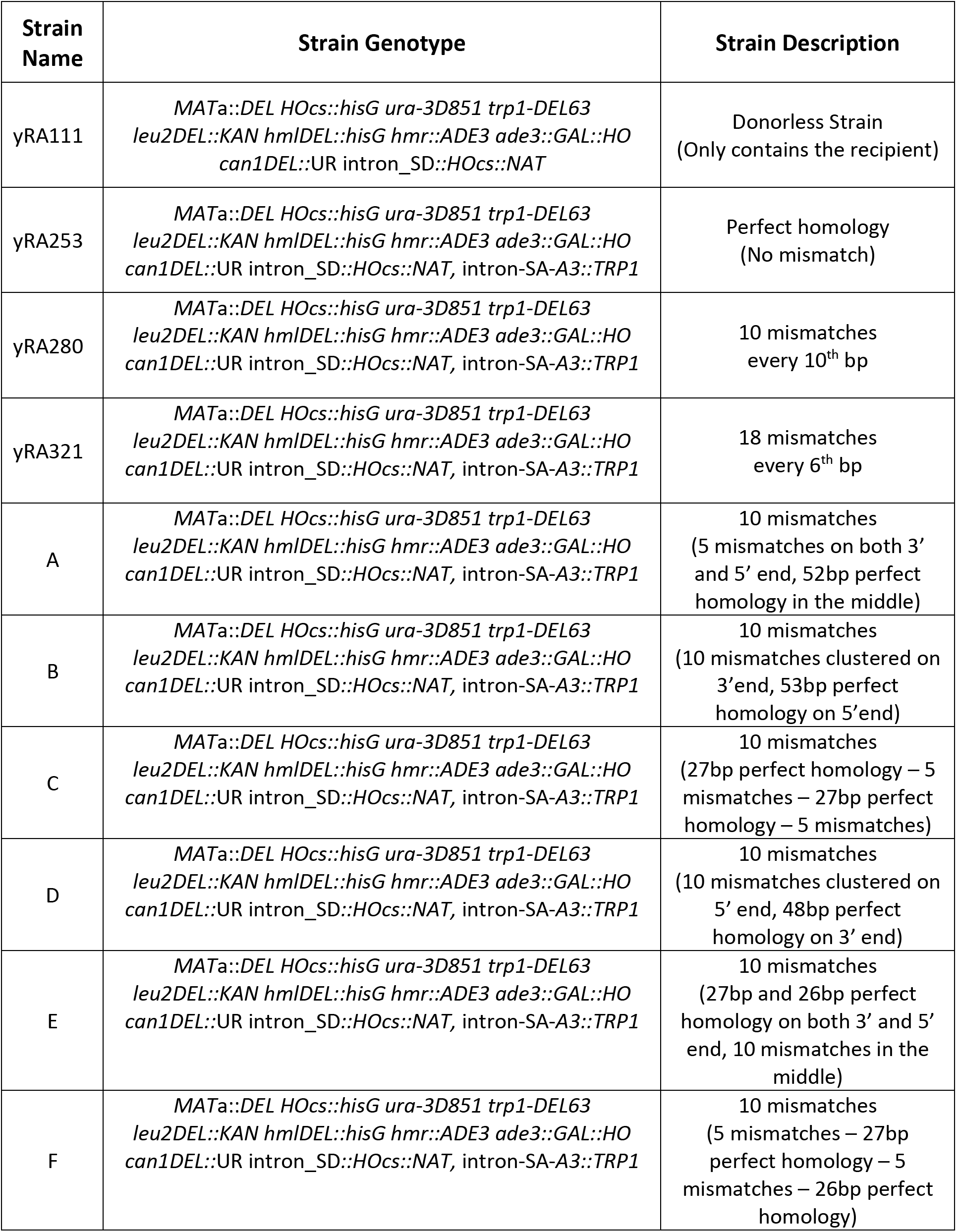
Yeast strains used in this study.

**Table 2.**
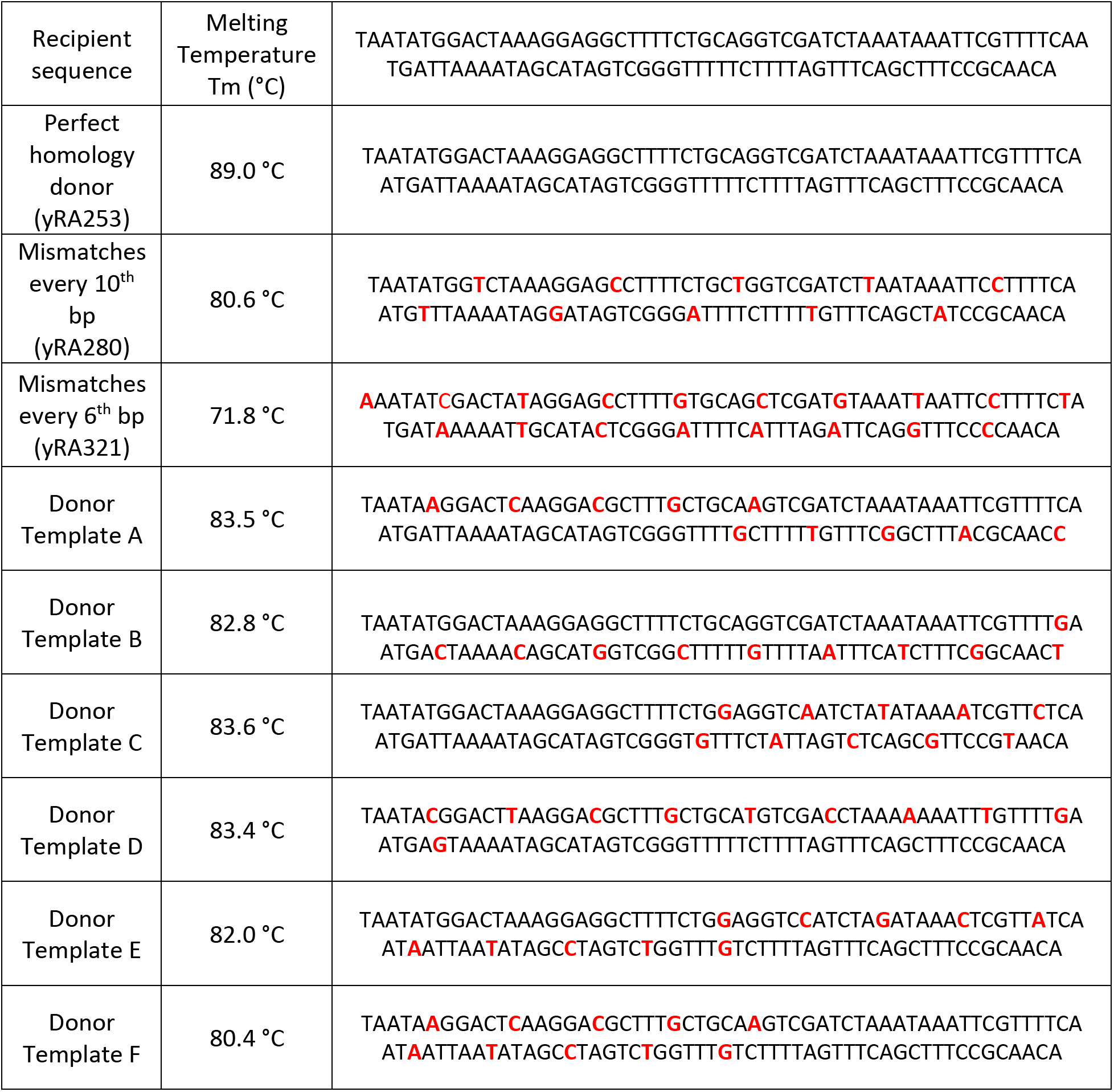
Recipient and donor sequences.

**Table 3.**
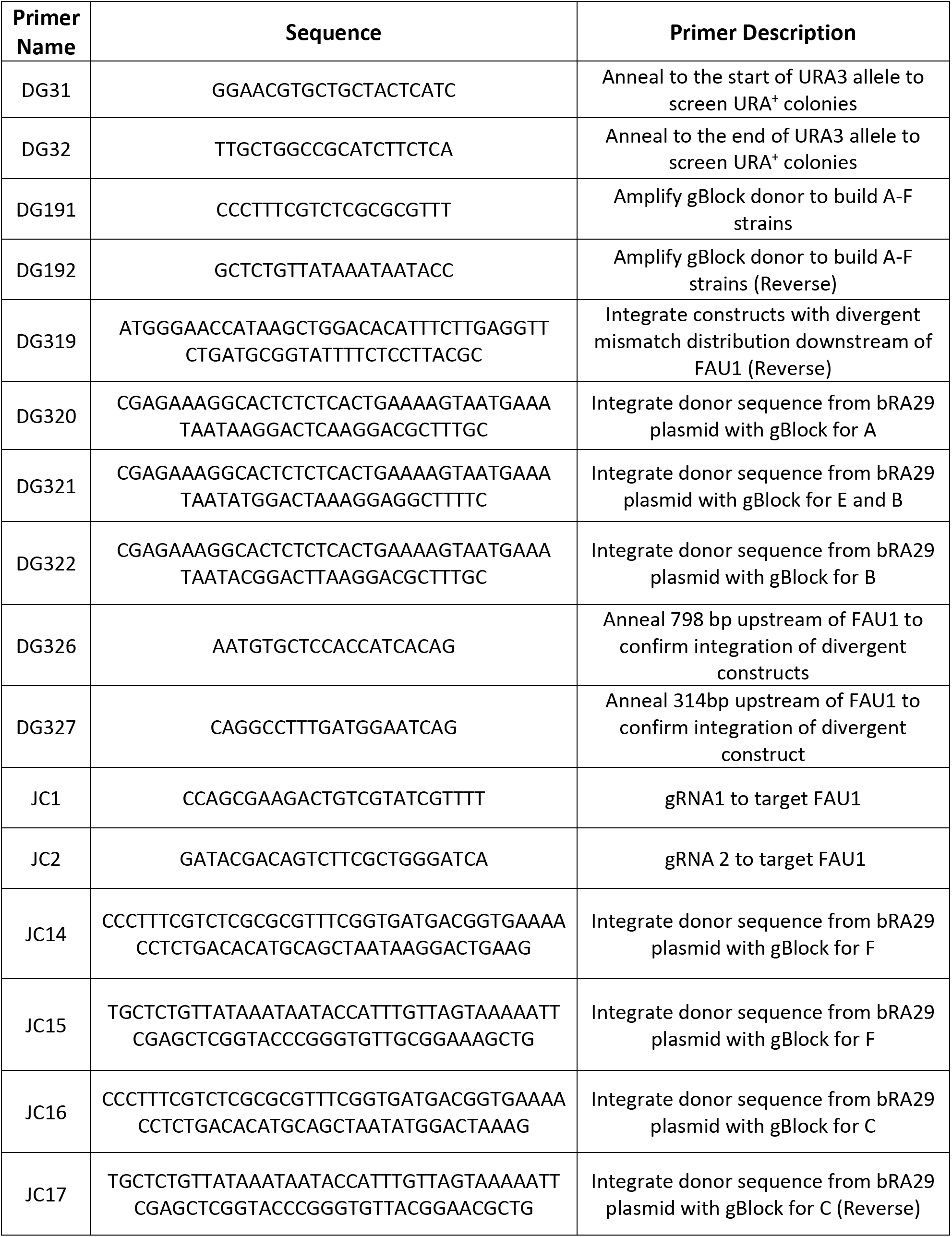
Primers used in this study.

**Table 4.** Plasmids used in this study.

| Plasmid Name | Plasmid Description |
| --- | --- |
| bRA25 | pRS314-based plasmid containing the 5' sequences from the *URA3* gene (UR), an artificial split-intron with the splice-donor site (5' SD) and the HO recognition site (HOcs) |
| bRA29 | pRS314-based (CEN4; TRP1) plasmid containing the 3' splice-acceptor of the intron, the 3' sequence from the *URA3* gene (A3), and the TRP 1 marker |

### BIR assay

Selected strains were grown on YEPD (1% yeast extract, 2% bactopeptone, 2% dextrose) + ClonNAT (100uL/mL final concentration) at 30°C. Individual colonies from strains were picked and serially diluted 1000-fold in double-distilled H_2_O. Serial dilutions were plated on YEPD to obtain the total number of cells and on YEP-galactose (YEP-Gal) to measure the recombination-dependent survivors after inducing HO endonuclease expression [36]. Cells were incubated at 30°C for 2-3 days. Cells on YEP-Gal were then replica plated to plates lacking uracil, to count colonies that survived the break via BIR, and to Nourseothricin (NAT) plates, to count colonies which survived the break via nonhomologous end-joining (NHEJ) that alters the HO cleavage site but retains the distal part of the left arm of chromosome V [3]. Ura^-^ NAT^+^ colonies, arising by nonhomologous end-joining, arose at a frequency of approximately 0.5% of the total number of colonies and are not reported for each assay.

### DNA sequence analyses of the break repair junctions

Representative Ura^+^ colonies that had repaired by BIR were confirmed by PCR, using primers amplifying the region at the start and end of the *URA3* gene using primers DG31 (GGAACGTGCTGCTACTCATC) and DG32 (TTGCTGGCCGCATCTTCTCA). PCR products were initially checked by gel electrophoresis and sent to GENEWIZ for Sanger sequencing. Individual PCR sequences were aligned with corresponding 108-bp donor templates and analyzed by DNA analyses software Serial Cloner 2.6.1 and Geneious Prime software.

### Statistical Analysis

GraphPad Prism 9 software was used to calculate statistical significance of data, based on Dunnett’s comparison of multiple samples versus a single control. Thermal stability, as reflected in melting temperature Tm (°C). of the base-pairing between the DSB 3’ and a complementary single strand of of the donor template was determined using the method of Markham and Zuker, 2005 (http://www.unafold.org/Dinamelt/applications/two-state-melting-hybridization.php) [37]. By using Geneious Prime’s DNA secondary structure fold viewer (https://assets.geneious.com/manual/2021.1/static/GeneiousManualse36.html), we monitored possible secondary structures of the donor templates. ΔG of all donor templates were calculated to determine the energy required to break the secondary structure.

## Acknowledgements

Funding: This work has been supported by the National Institute of Health grants R35 GM127029 to J.E.H D.N.G. has been supported by NIGMS Genetics Training Grant T32GM007122 (https://www.nigms.nih.gov/) and by the National Science Foundation Graduate Research Fellowship Program under grant 1744555 (https://www.nsfgrfp.org/). Special thanks to Dr, Michael Lichten (National Cancer Institute Center for Cancer Research), Dr. Stephen Levene (University of Texas at Dallas), Dr. Eric Greene (Columbia University), Dr. Myron Goodman and Dr. Chiho Mak (University of Southern California) for discussions about this research.

---

**Figure S1.**
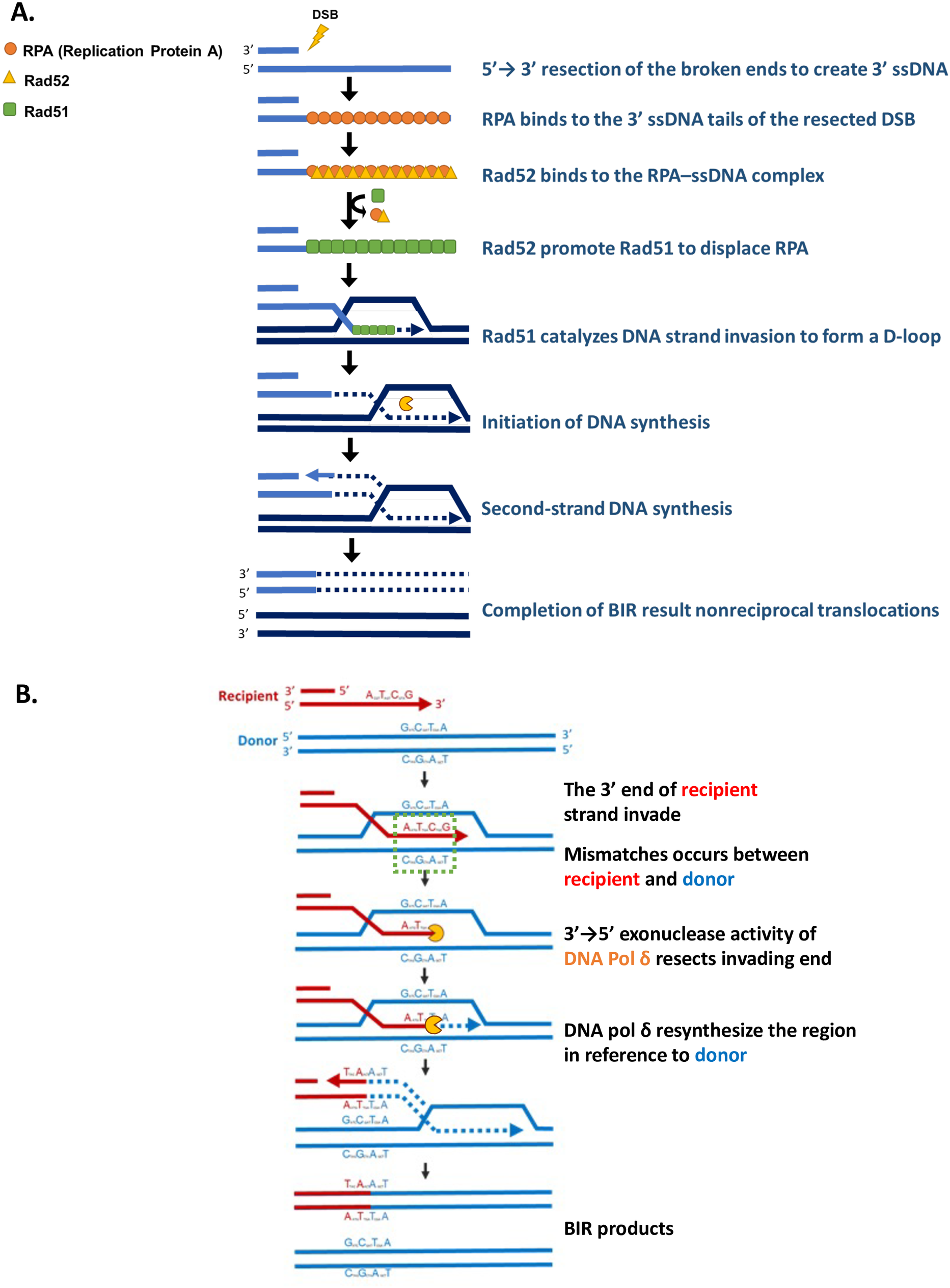
Rad51-mediated Break-Induced Replication. **A.** Mechanism of Rad51-dependent BIR **B.** Mismatch incorporation of heteroduplex DNA formation during BIR. Once a DSB is created, a broken end of DSB will be resected by 5’→3’ exonuclease to generate 3’ single-stranded DNA (ssDNA) which interact with several recombination proteins to carry out homology search and strand invasion. Since both donor and recipient sequence share a 108-bp homology, the broken and resected end of recipient sequence (indicated in red) will find this homology and initiate strand invasion into donor sequence (indicated in blue). Mismatches are created in the heteroduplex region during strand invasion and polymerase δ get recruited to perform its proofreading 3’→5’ exonuclease activity. DNA Polymerase δ “backs up” into the heteroduplex region and resynthesizes the region in reference to donor templates to incorporate mismatches. BIR completes and results in BIR products with mismatches close to the 3’ end frequently being incorporated.

**Figure S2.**
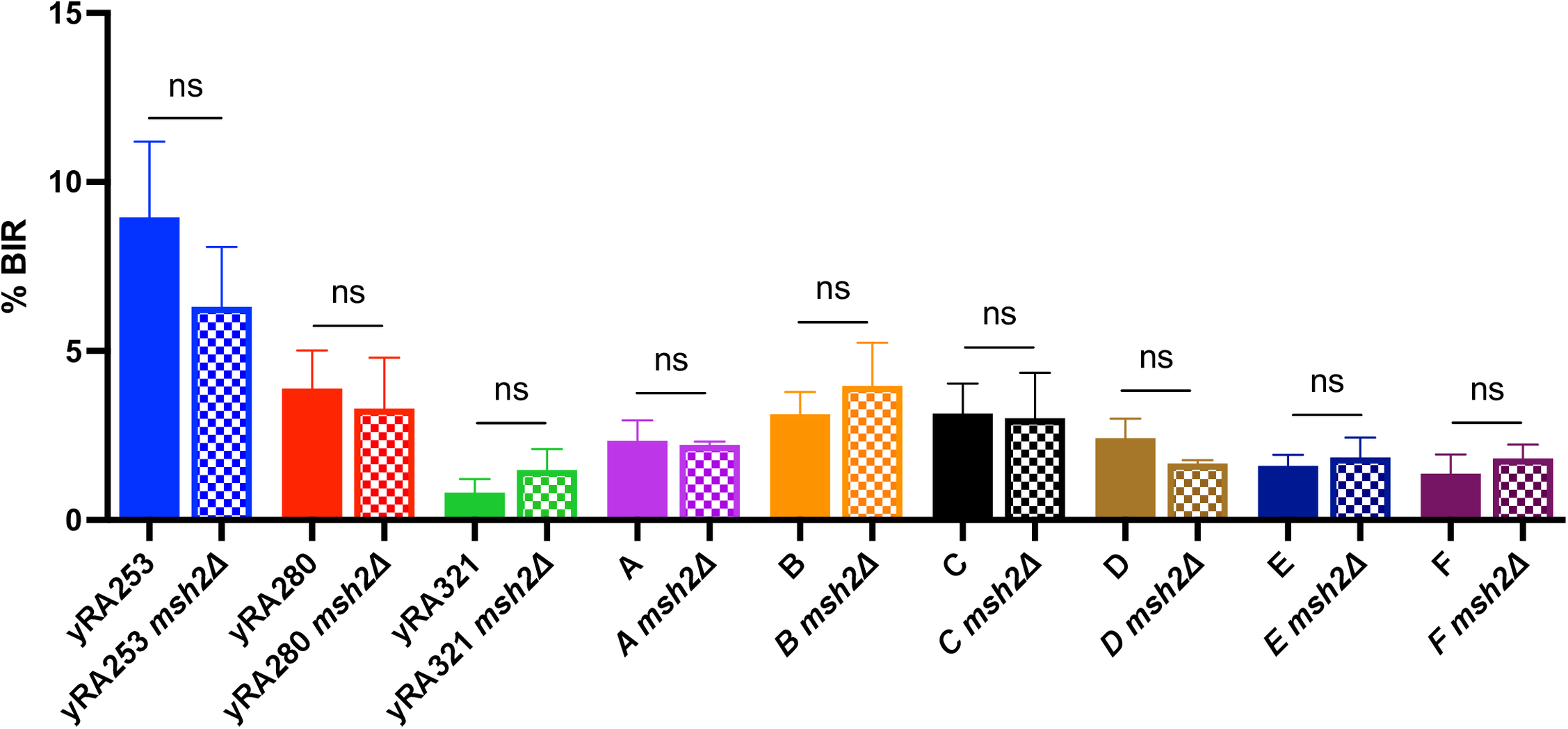
The effect of *MSH2* mutants on the repair efficiency. Wild type and *msh2*Δ derivatives of each donor template were measured as described in Fig. 1. Statistical significance of the differences for each donor/*msh2*Δ pair was determined using an unpaired t-test with Welch’s correction. Error bars refer to standard deviation. Each measurement is based on a minimum of three experiments.

